# High life history diversity within a single genus of phytoplankton viruses

**DOI:** 10.1101/2022.03.13.484168

**Authors:** Eva J.P. Lievens, Irina V. Agarkova, David D. Dunigan, James L. Van Etten, Lutz Becks

## Abstract

Phytoplankton viruses are key players in aquatic ecosystems, where they control algal populations and affect nutrient flow. The ecological impact and evolution of these viruses can be understood by studying their life history traits, but very little is known about the life history diversity of related viruses. We quantified the life cycle of 34 strains in the genus *Chlorovirus*, which infects freshwater green algae. All chloroviral life history traits varied 5-to 75-fold across strains, in some cases rivaling the known trait range for all phytoplankton viruses. The trait variation affected viral growth rates but was not detectably constrained by life history trade-offs. This study represents the most in-depth characterization of algal viruses to date and raises the question whether all phytoplankton virus genera are equally diverse.

## Introduction

Phytoplankton viruses play an important role in aquatic ecosystems through their ability to lyse photosynthesizing microbes. An effective way to understand this role is to study the traits that determine viral fitness: their life history or ‘performance’ traits [1]. For lytic viruses, classic life history traits include lysis time, burst size, and mortality rate. These traits shape viral effects on host populations, and can thus be used to parameterize ecological models that assess viral impacts [2].

Life history traits are also used to look for overarching constraints and patterns in viral evolution [3–6]. The life histories of phytoplankton viruses are highly diverse, consistent with the wide range of genome types, morphologies, host ranges, and abiotic environments represented in this functional group [5–7]. However, very little is known about the diversity of phytoplankton viruses at the genus and species level. This limits our ability to assess the variation in ecological impact of similar viruses, and to assess the ecological consequences of viral population evolution.

We studied life history diversity in chloroviruses (family *Phycodnaviridae*, genus *Chlorovirus*), a particularly well-characterized genus of phytoplankton viruses. These lytic dsDNA viruses i) infect unicellular green algae in freshwater environments [8], ii) have a well-described life cycle [9], and iii) contain over 130 isolated and sequenced strains [10]. We characterized the life histories of 34 strains belonging to the subgenera *Alphachlorovirus* (species I, II, and V) and *Gammachlorovirus* (species not yet assigned), which infect *Chlorella variabilis* and *Chlorella heliozoae* respectively [10]. We then compared the *Chlorovirus* trait diversity with that of other phytoplankton viruses and explored its consequences for chloroviral ecology and evolution.

## Results & Discussion

We used previously described modified one-step growth and survival assays to quantify the chlorovirus’ life history as follows (Fig. 1A) [9]. Chloroviruses first adsorb to the host at a rate determined by the adsorption constant *k*, then depolarize the host’s plasma membrane with depolarization probability *d*. Depolarization is followed by viral genome entry, replication, and release of progeny virions (virus particles) through lysis; the probability that these steps are successful is the release probability *r*. Release timing is described by the mean ± SD of lysis time *μ*_*l*_ ± *σ*_*l*_, and the number of progeny virions is the burst size per depolarized cell *b*_*d*_ or per release *b*_*r*_. The overall probability that a virion can complete this cycle (i.e. that a virion is infectious) is the specific infectivity *s*. In the absence of available hosts, most infectious virions decay following mortality rate *m*, but a persistent fraction *p* resists decay. Importantly, the depolarization probability, release probability, and persistent fraction were inaccessible prior to the development of these assays [9].

**Fig. 1.**
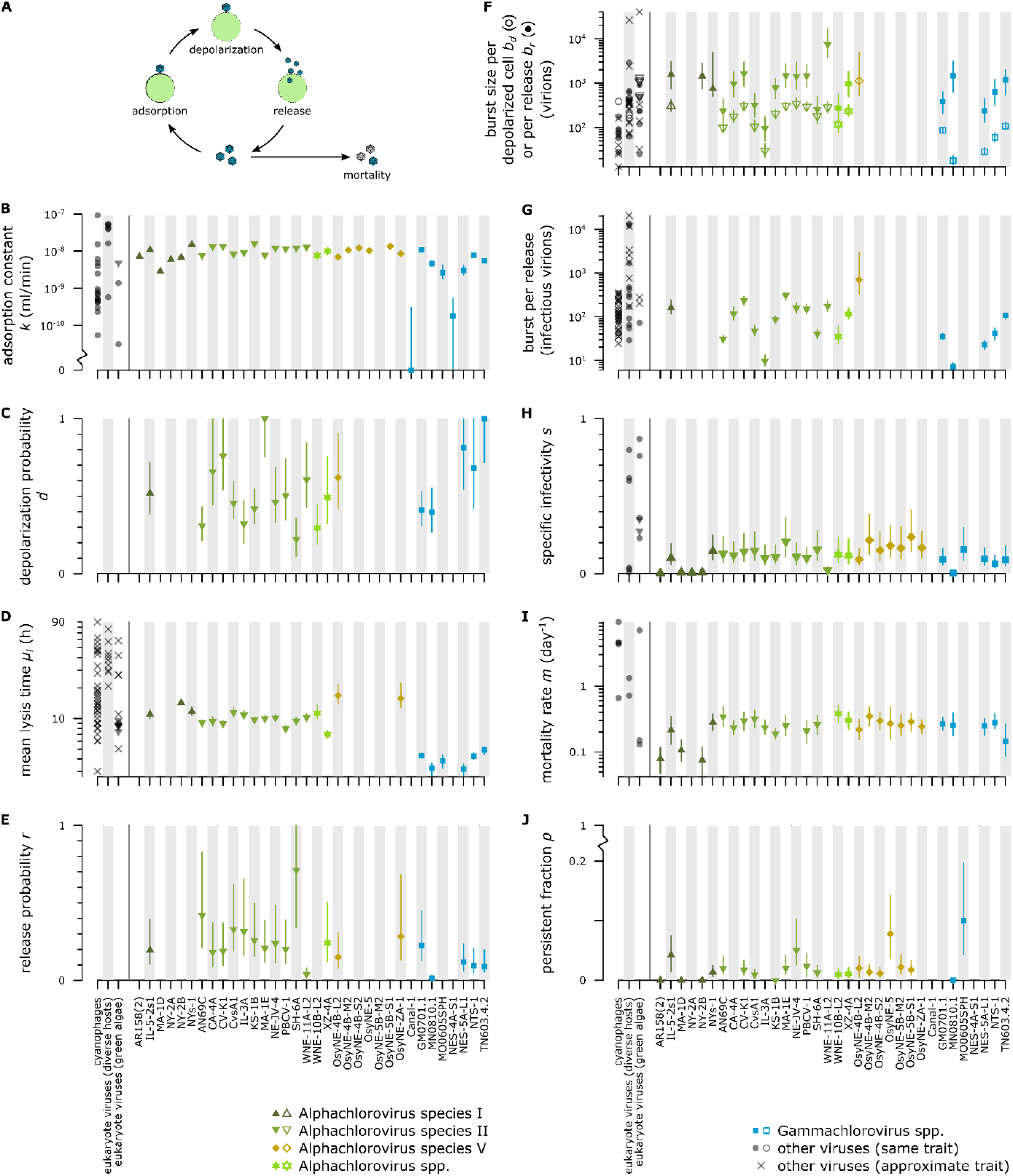
Life history of the chloroviruses. **A. Overview of the chlorovirus life cycle**. Each step is associated with one or more life history traits. **B-J. Life history traits**. Chlorovirus data is shown as trait estimates with 95% CIs; blanks indicate poorly resolved traits. SD of lysis time is not shown as it was tightly correlated with mean lysis time. Chlorovirus data was compared to published values for phytoplankton viruses (dark grey); published values were “approximate” if we manually inferred the mean lysis time, or if it was unclear whether burst size was measured per lysed or infected (≈ depolarized) cell. See Supporting Information for details and references.

We found that the chlorovirus traits were remarkably diverse, with trait estimates varying 5-to 75-fold between strains (Fig. 1B-J). Moreover, *Chlorovirus* adsorption constants and burst sizes spanned most of the known trait range for phytoplankton viruses (Fig. 1B,F,G). This is particularly striking because the known trait range was obtained in a variety of host species and environments, while the chlorovirus traits were not [though all environments were permissive, following 5,6]. In contrast, *Chlorovirus* lysis time, specific infectivity, and mortality rate had more restricted trait ranges compared to other phytoplankton viruses (Fig. 1D,H,I). This restriction might be a proximate consequence of shared host physiology (e.g. high growth rate causing fast lysis [5]) or an ultimate consequence of shared ecological conditions (e.g. slow mortality to cope with unpredictable host availability).

To explore how trait variation affects population dynamics, we measured viral growth in short-term assays. Observed growth rates correlated with growth rates predicted from our estimates of specific infectivity, adsorption constant, mean lysis time, and burst size per release (ρ = 0.51, *p* = 0.02; Fig. 2A). However, regressing the observed growth rate onto these traits showed that only specific infectivity had a strong effect (present in all models with ΔAIC ≤ 2, *n* = 21 strains; Fig. 2A). Thus, trait variation affected the chlorovirus’ ecological dynamics, but not all traits were equally relevant. Future work should test whether relevance is context-dependent, e.g. whether adsorption constant is more important when host densities are lower.

**Fig. 2.**
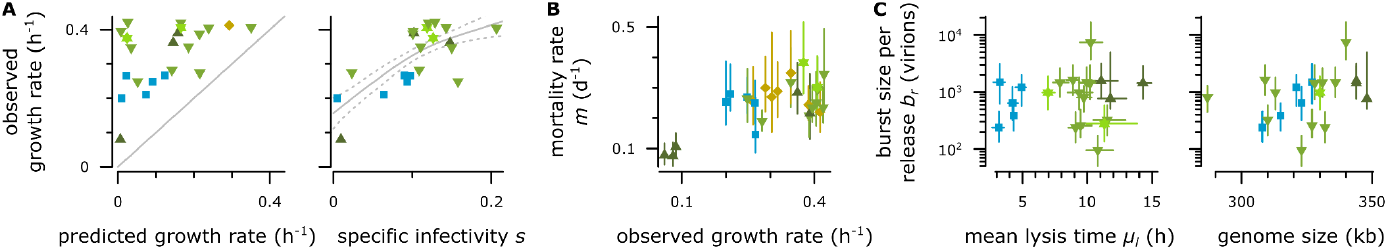
Viral growth and potential trade-offs. **A. Trait basis of viral growth**. Left: Observed growth rate was compared with the growth rate predicted from the adsorption constant, mean lysis time, specific infectivity, and burst size (gray: 1:1 line). Right: Of the aforementioned traits, only specific infectivity had a detectable effect (gray: prediction ± SE of the relevant regression model). **B-C. Tested trait correlations**. Analyses were weighted by 95% CI breadth. Symbols and colors as in Fig. 1.

Finally, we looked for evidence that chlorovirus evolution is constrained by trade-offs between life history traits. We tested for correlations consistent with three trade-offs that are hypothesized to affect lytic viruses: growth rate vs. mortality rate [3,4], burst size vs. lysis time [4], and burst size vs. genome size [6]. None of the hypothesized trade-offs were supported within *Alphachlorovirus* species I, II, or V, nor in the *Gammachlorovirus* subgenus (Fig. 2B,C). Indeed, genome size and burst size were positively correlated in Gammachlorovirus (*ρ* = 0.98, *p* = 0.03; Fig. 2C). The trade-offs may have been obscured by other differences between the strains [11]; alternately, they may be mitigated by additional functions in the large chlorovirus genome [6].

In summary, we found that the *Chlorovirus* genus contains high diversity in traits with direct effects on fitness. Thus we expect different chloroviruses to have very different impacts on host populations, both in the short term (Fig. 2A) and the long term (Fig. 1I-J), and we expect chlorovirus populations to respond quickly to selection [cf. 12]. The scale of *Chlorovirus* trait diversity is remarkable when compared to that of other phytoplankton viruses, in particular for burst size (Fig. 1F-G). If other virus genera are similarly diverse, this implies that known viruses may not be representative of their relatives. This would complicate the bottom-up prediction of host-virus dynamics, which would be sensitive to the specific viruses measured and to evolutionary shifts in the population, as well as the inference of overarching life history strategies. Exploring the phenotypic diversity of phytoplankton viruses at the genus and species level is therefore an essential next step to understanding their function in aquatic ecosystems.

## Methods

We used 34 chlorovirus strains belonging to the subgenera *Alphachlorovirus* and *Gammachlorovirus*. Strains were phenotyped in their type hosts [10]: alphachloroviruses in *C. variabilis* NC64A (species I, II, and unknown; *n* = 6, 11, and 2) or *C. variabilis* Syngen 2-3 (species V; *n* = 7), and gammachloroviruses in *C. heliozoae* SAG 3.83 (species unknown; *n* = 8). Algae were in a late exponential growth phase.

We measured the chlorovirus’ life history traits using modified one-step growth and survival assays [9]. These methods use statistical modeling to infer life history traits from population-level measurements; they generate a point estimate and 95% confidence interval for each trait. Poorly resolved estimates were excluded from further analysis (blanks in Fig. 1B-J).

The virus’ observed growth rate was measured over a 24 h period in liquid culture (replicated twice). The observed growth rate was calculated as *ln*(final/initial virion concentration)/24h. The predicted growth rate was calculated as De Paepe & Taddei’s multiplication rate [3] with an additional adsorption term (see Supporting Information). We used Spearman’s rank correlation to compare the two measures, and shape-constrained additive models to regress the observed growth rate onto the component traits [13].

To test for signals of trade-offs, we used Spearman’s rank correlations to compare growth rate vs. mortality rate [3,4], burst size vs. lysis time [4], and burst size vs. genome size [6]. Correlations were weighted by the precision of each estimate [1/√(product of the CI breadth of trait 1 and trait 2), 14]. Each correlation was tested within species where *n* ≥ 5, with subgenus *Gammachlorovirus* treated as one species.

Details on culturing conditions, virus stocks, assays, and analyses can be found in the supporting information. Data and code are available at https://doi.org/10.5281/zenodo.6573769 and https://doi.org/10.5281/zenodo.13999011.

## Acknowledgements

We thank J. Clot, R. Hermann, M. Duffy, and the University of Konstanz flow cytometry centre (FlowKon). E.J.P.L. acknowledges the University of Konstanz Young Scholar Fund.

## Notes

https://www.protocols.io/view/virion-quantification-by-flow-cytometry-without-fi-6qpvr6q93vmk/v1

https://www.protocols.io/view/modified-one-step-growth-mosg-assay-4r3l2op9pv1y/v3

https://www.protocols.io/view/modified-persistence-mp-assay-81wgb6knylpk/v3

### Competing Interest Statement

The authors have declared no competing interest.

### Summary of Updates

Update to reflect that the previous version was split into two manuscripts. The methods half was published at https://doi.org/10.1128/aem.01659-23; the diversity half is updated here.

