## supporting methods for "High life history diversity within a single genus of phytoplankton viruses"

### Note on the supporting methods

In this study, we used recently-developed methods to characterize 34 chlorovirus strains. Four of these strains were also included in Lievens et al. [1]*, where they illustrated the development of the experimental and analytical methods. As some aspects of the analytical workflow have to be run on the entire dataset**, the raw data and kinetic models for all 34 strains were made available at https://doi.org/10.5281/zenodo.6573769 upon publication of Lievens et al. [1]. Curated data and code for the current analyses are available at https://doi.org/10.5281/zenodo.13999011 .

**Alphachlorovirus* strains AN69C, CV-K1, KS-1B, and PBCV-1.

**Specifically the decision to simplify the kinetic models, see Lievens et al. [1] Figs. S8 and S10.

### Supporting methods

*Viruses, algae, and experimental conditions*

Unless otherwise specified, assays took place at 20°C and under constant light. Algal growth and virus incubations were done on an orbital shaker with diameter 10 mm and frequency 120 rpm. Any 4°C storage was also dark storage.

We phenotyped chloroviruses belonging to the subgenera *Alpha*- and *Gammachlorovirus* (Table S1). Viruses belonging to *Alphachlorovirus* species I and II infect the ciliate endosymbionts *Chlorella variabilis* NC64A and *C. variabilis* Syngen 2-3. Their type host is *C. variabilis* NC64A. Viruses belonging to *Alphachlorovirus* species V have diverse host ranges; we tested a set of strains that can only replicate in *C. variabilis* Syngen 2-3. Viruses belonging to the subgenus *Gammachlorovirus*, which has not yet been divided into species, infect the heliozoon endosymbiont *Chlorella heliozoae* SAG 3.83. All of the described procedures took place in the type hosts.

**Table S1. List of viral strains.** All isolates/strains can be found at NCBI_BioProject PRJNA1154233. Species assignments following Carvalho et al. [2].

| Subgenus | Species | Strains | Type host |
|---|---|---|---|
| *Alphachlorovirus* | I | AR158(2)*, IL-5-2s1, MA-1D, NY-2A, NY-2B, NYs-1 | NC64A |
| | II | AN69C, CA-4A, CV-K1 (also called CviKI), CvsA1, IL-3A, KS-1B, MA-1E, NE-JV-4, PBCV-1, SH-6A, WNE-11A-L2 (also called WNE-11A-L1) | NC64A |
| | V | OSyNE-4B-L2, OSyNE-4B-M2, OSyNE-4B-S2, OSyNE-5, OSyNE-5B-M2, OSyNE-5B-S1, OSyNE-ZA-1 | Syngen 2-3 |
| | unknown | WNE-10B-L2, XZ-4A | NC64A |
| *Gammachlorovirus* | unknown | Canal-1, GM0701.1, MN0810.1, MO0605SPH, NES-4A-S1, NES-5A-L1, NTS-1, TN603.4.2 | SAG 3.83 |

*Suffix (2) used to distinguish this lysate from another AR158 lysate in our collection.

The chlorovirus strains were isolated from natural ponds or streams around the world between 1981 and 2017, and have been maintained as part of a stock collection since then. Virus suspensions were stored in lysate form at 4°C. Before starting the assays, we refreshed and amplified the virus stocks: 0.5 ml of the lysate was inoculated into 10 ml of $2 \times 10^6$ algae/ml suspension, incubated until the algae lysed (24 h or 48 h), and filtered through 0.2 µm. The resulting filtrates were stored at 4°C and the concentration of virus particles (virions) was measured by flow cytometry [1,3]. The filtrates with insufficient virion concentrations were given one more round of this amplification treatment.

The algal strains NC64A, Syngen 2-3, and SAG 3.83 were stored on agar slants at 4°C and inoculated into liquid medium before use. We used a modified version of Bold's Basal Medium [BBM, 4], with ammonium chloride substituted for sodium nitrate as a nitrogen source and double the concentration of trace element solution 4 [first used by 5]. We keep the abbreviation BBM in order to distinguish our medium from the enriched "MBBM" typically used in this model system [e.g. 6]. Algae used in the assays were in late exponential phase (~$2 \times 10^6$ algae/ml in this medium).

*Measurement of life history traits*

We measured the chlorovirus'  life history traits using modified one-step growth (mOSG) and modified survival (mS) assays, as described in Lievens et al. [1] and on protocols.io [7,8]. Briefly: The mOSG assay tracked virion concentrations over the course of a single replication cycle. Importantly, replicate infections were initiated at varying virion:host ratios. A kinetic model was fit to the virion concentrations and used to estimate point estimates and 95% CIs for most of the trait shaping viral reproduction (Table S2). The mS assay tracked the decline of infectious virions in the environment when exposed to 20°C and constant light. A kinetic model was fit to the infectious virion concentrations and used to estimate point estimates and 95% CIs for the specific infectivity and survival traits (Table S2). By comparing the mOSG and mS assays, we were able to estimate point estimates and 95% CIs for the two remaining reproduction traits (Table S2).

**Table S2. Traits estimated in the mOSG and mS assays.**

| Life cycle phase | Symbol | Trait | Description | Assay |
|---|---|---|---|---|
| Reproduction | $k$ | adsorption constant | determines the rate at which virions adsorb to host cells | mOSG |
| | $d$ | depolarization probability | probability that an adsorbed virion is capable of depolarizing a host cell (can be quantified through the variation in initial virion:host ratio) | mOSG |
| | $\mu_l$ | mean lysis time | mean lysis time, assuming all release events produce the same number of virions | mOSG |
| | $\sigma_l$ | standard deviation of lysis time | standard deviation of lysis time, assuming all release events produce the same number of virions | mOSG |

| | | | | |
|---|---|---|---|---|
| | $r$ | release probability | probability that a depolarized cell releases infectious virions | mOSG & mS |
| | $b_d$ | burst size per depolarized cell | average number of virions produced by a depolarized cell | mOSG |
| | $b_r$ | burst size per release | average number of virions produced per release event | mOSG & mS |
| | $s$ | specific infectivity | initial proportion of infectious virions (equivalent to the probability that a virion is capable of adsorbing, depolarizing, and releasing infectious progeny) | mS |
| Survival | $m$ | mortality rate | rate at which nonpersistent infectious virions became noninfectious | mS |
| | $p$ | persistent fraction | proportion of persistent infectious virions, i.e. of infectious virions that resisted decay for the duration of the assay (4 weeks) | mS |

After fitting the kinetic models, we evaluated the fits and trait estimates. We excluded trait estimates from incomplete assays ($d$, $\mu_l$, $\sigma_l$, $b_d$, $r$, and $b_r$ for strains that did not finish growing in the mOSG assay; $m$ and $p$ for strains that did not decay in the mS assay; all traits for assays that failed), estimates derived from poorly fitting models, and estimates with uninformatively broad CIs compared to the plausible phenotypic range (criteria: CI > 0.67 for $d$, $s$, and $r$; CI spanning more than an order of magnitude for $b_d$, $p$, and $b_r$; CI overlapping with the upper fitting constraint for $k$, $\mu_l$, $\sigma_l$, and $m$). Details on exclusion are provided in Table S3.

**Table S3. Trait estimates excluded from further analysis.**

| Reason | Assay | Traits excluded | Strains |
|---|---|---|---|
| Incomplete assays | mOSG | $k$, $d$, $\mu_l$, $\sigma_l$, $b_d$, $r$, $b$ | OSyNE-5B-M2 |
| | | $d$, $\mu_l$, $\sigma_l$, $b_d$, $r$, $b_r$ | AR158(2), Canal-1, MA-1D, NES-4A-S1, NY-2A, OSyNE-4B-M2, OSyNE-4B-S2, OSyNE-5, OSyNE-5B-S1 |
| | mS | $s$, $m$, $p$ | Canal-1, NES-4A-S1 |
| | | $m$, $p$ | NY-2A |
| Poor model fit | mOSG | $d$, $b_d$, $r$, $b_r$ | MO0605SPH |
| Uninformative CIs | mOSG | $d$ | NY-2B, NYs-1, OSyNE-ZA-1 |
| | | $b_d$ | NY-2B, NYs-1, OSyNE-ZA-1 |
| | | $r$ | NY-2B, NYs-1, WNE-10-L2 |
| | | $b_r$ | OSyNE-4B-L2, OSyNE-ZA-1 |
| | mS | $m$ | MO0605SPH, NE-JV-4, WNE-11A-L2 |
| | | $p$ | CA-4A, IL-3A, GM0701.1, NES-5A-L1, NTS-1, OSyNE-ZA-1, TN603.4.2, WNE-11A-L2 |

A second evaluation step is to examine the effect of correlations between model parameters [compensating effects, 1]. These correlations are caused by random noise and are inherent to the model fitting. For example, if the initial concentration of infectious virions in the mS assay is estimated to be slightly higher/lower than the true value, then model fitting can lead to an over-/underestimate of both the specific infectivity and the mortality rate. Lievens et al. found that correlations occur between parameters $d$ & $b_d$, $\mu_l$ & $\sigma_l$, $\mu_l$ & $b_d$, $\sigma_l$ & $b_d$, $s$ & $p$, and $s$ & $m$ [Fig. S4 in 1]. A visual assessment of these effects in the current dataset is presented in Fig. S1.

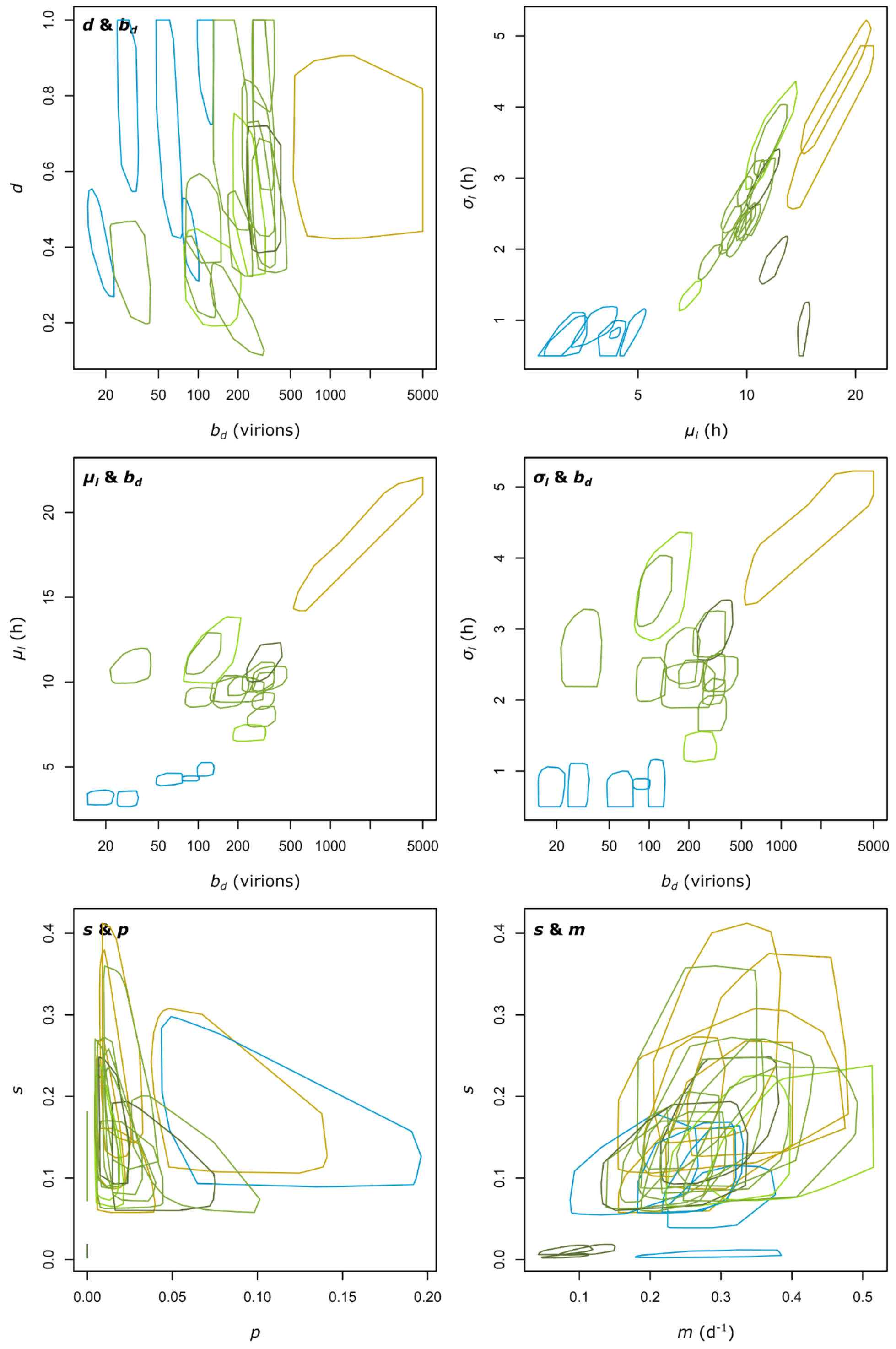


**Fig. S1. Effect of correlations between model parameters arising from the fitting process.** To look for confounding effects of the correlations between parameters $d$ & $b_d$, $\mu_l$ & $\sigma_l$, $\mu_l$ & $b_d$, $\sigma_l$ & $b_d$, $s$ & $p$, and $s$ & $m$, we compared the bootstrapped CIs to the overall phenotype space. Each panel shows a parameter combination, and each polygon represents one viral strain. Colors as in Fig. 1: green shades indicate *Alphachlorovirus* species I, II, and unknown (tested in *C. variabilis* NC64A), gold indicates *Alphachlorovirus* species V (tested in *C. variabilis* Syngen 2-3), and blue indicates *Gammachlorovirus* strains.. Polygons outline the bootstrap values that fell within the 95 % CIs. Example interpretation 1: there is little overlap between polygons for $\mu_l$ and $b_d$, so parameter correlations do not need to be considered further. Example interpretation 2: the overlapping polygons for *s* and *p* indicate that correlations caused by model fitting cannot be distinguished from true phenotypic correlations.

The biological assumptions underlying the mOSG and mS assays are discussed in detail in Lievens et al. [1]. Here we highlight two assumptions that are particularly relevant to the current dataset. First, we assumed that all virions were capable of adsorbing to host cells. If a non-negligible proportion of virions could not adsorb (in all or some strains), then we have underestimated the adsorption constant and release probability and overestimated the burst size per release (in all or some strains). Second, we assumed that the most probable number-like approach used in the mS assay accurately measured the infectiousness of virus suspensions. Specifically, we assumed that all algal cultures inoculated with ≥ 1 infectious virion lysed within 4 days. This may not be the case for slow-growing strains (e.g. MA-1D, NY-2B), which would lead to underestimates of the specific infectivity and release probability and overestimates of the burst size per release.

All analyses were performed in R [9].

*Comparison with published trait data*

To obtain a range of trait data for phytoplankton viruses, we combined data from published reviews with updates from a systematic literature search. For optimal comparability, we only retained phenotypes obtained under permissive conditions [growth traits, following 10,11] or well-lit conditions (mortality rate). Where necessary, data was extracted manually using WebPlotDigitizer version 4 [12]. For the precise search procedure, search terms, extracted data, and references, see Supporting Methods File 2.

- For the adsorption constant, we used the data collected by Edwards et al. [11] as a basis and searched Web of Science for estimates published since 2020.
- For lysis time and burst size, we used the data collected by Edwards & Steward [10] as a basis and searched Web of Science for estimates published since 2018. For any papers that reported the earliest lysis time (latent period) or latest lysis time, we extracted the approximate mean lysis time (the half-way point of the virus rise curve or of the host lysis curve). We noted whether burst size was measured in virions or infectious virions. We also noted whether burst size was calculated per infected cell (most similar to our burst size per

depolarized cell) or per lysed cell (most similar to our burst size per release); if this was unclear we scored the trait as approximate.

- For specific infectivity, we searched Web of Science for published estimates.
- For mortality rate, we used the data collected by Mojica & Brussaard [13] as a basis and searched Web of Science for estimates published since 2014.

*Growth assay*

We used the refreshed filtrates prepared at the start of the mS assay to measure viral growth. Viruses and algae were combined in a 96-well culture plate (2 wells per strain) to a final volume of 0.2 ml containing $1 \times 10^6$ algae/ml and 2,500 or 25,000 virions/ml (virion:host ratio 0.0025 or 0.025). The suspensions were incubated for 24 h. After 24 h, suspensions were centrifuged for 15 min at 2000 g to separate algae and viruses, and the virion concentration of the supernatants was measured by flow cytometry [1,3]. We calculated the observed growth rate as *ln*(mean of the final/initial virion concentration for the 2 replicates)/24.

The predicted growth rate was calculated as De Paepe & Taddei's multiplication rate $s*b_r*\mu_l^{-1}$ [14] with an additional adsorption term: $(1 - e^{-k*A_g}) * s * b_r * \mu_l^{-1}$ or equivalently $(1 - e^{-k*A_g}) * d * b_d * \mu_l^{-1}$. Here $A_g$ is the algal concentration during the growth assay, i.e. $1 \times 10^6$ algae/ml.

The observed and predicted growth rate were compared using Spearman's rank correlation. To examine whether all component traits contributed to the variation in observed growth rate, we regressed the observed growth rate onto the adsorption constant *k*, specific infectivity *s*, mean lysis time $\mu_l$, and *ln*-transformed burst size per release $b_r$. We allowed the observed growth to be a nonlinear function of the trait values, but we forced the functions to be mechanistically sensible: adsorption, specific infectivity, and burst size were constrained to positive monotonic effects, while lysis time was constrained to a negative monotonic effect [shape-constrained additive models, package "scam" version 1.2-17, 15]. We compared models with all additive trait combinations using the Akaike information criterion [AIC, 16]; traits present in all models with $\Delta$AIC ≤ 2 were considered to be predictive. Analyses were performed in R [9].

*Signals of trade-offs*

Finally, we looked for signals of three trade-offs that are commonly hypothesized to affect lytic viruses: growth rate vs. mortality rate [17,18], burst size vs. lysis time [18], and burst size vs. genome size [19]. To avoid phylogenetic artifacts we tested within virus species, with subgenus

*Gammachlorovirus* treated as one species. We did not test correlations if $n < 5$. Correlations were tested using Spearman's rank correlations, weighted by the precision of each estimate [1/√(product of the CI breadth of trait 1 and trait 2), package 'weights', 20]. Holm's method was used to correct for multiple testing [21]. Analyses were performed in R [9].

*References*

[12] Rohatgi A. WebPlotDigitizer: Version 4.8. https://apps.automeris.io/wpd4/

[20] Pasek J. weights: Weighting and Weighted Statistics, version 1.0.4 2021.

[21] Holm S. A simple sequentially rejective multiple test procedure. Scand J Stat 1979;6:65–70.
